# Effects of age and sex on diet and activities of immature reintroduced western lowland gorillas

**DOI:** 10.64898/2026.02.02.703196

**Authors:** Anna Cryer, Tony King, Nicaise Ngoulou, Elke Boyen, Sander Muilerman-Rodrigo, Julia Lehmann

**Author notes:** Corresponding Author: Anna Cryer.

## Abstract

The juvenile life stage is crucial in primates, yet the behavior and diet of juveniles is an understudied area of primatology. Compared with adults, considerably less is known about wild juvenile primate development, particularly that of western lowland gorillas (*Gorilla gorilla gorilla*). While sex differences in diet and time budgets are well studied in adults, work remains to be done on how age and sex influence juvenile behavior. Here, we use data from nine immature reintroduced western lowland gorillas to fill some of this knowledge gap. We found that the immature gorillas spent most time feeding, followed by resting. Younger juveniles spent less time resting than older individuals, instead spending more time in self-play compared with older juveniles and more time in locomotion than subadults. The group had a varied diet as would be expected for western lowland gorillas; predominantly eating stems, flowers/leaves and fruit, with subadults eating more stems compared to younger individuals. Sex was found to have little influence on either behavior or diet. Additionally, behavioral and dietary diversity were calculated in order to describe the diversity of immature western lowland gorilla behavior. There were no age or sex differences found among this group of individuals, suggesting behavioral repertoire and diet may be influenced by those in their social group. The wider aim of this study is to provide insights into immature western lowland gorilla behavior and diet in the wild while also contributing to understanding of the post-release period for rehabilitated primates.

## Introduction

Primates are unique among mammals with an extended juvenile and subadult period (Lonsdorf, 2017) and this period is central to development. Juvenile nonhuman primates (hereafter primates) are difficult to study in the wild due to their size, unpredictable movement and complexities in the identification of individuals (Barale et al., 2015). Many studies consider juveniles as a homogenous group, not taking the sex or age of individuals into consideration (e.g. Watts, 1988; Cooper and Bernstein, 2000; Sueur et al., 2011; Liu et al., 2016). Therefore, despite the known importance of the juvenile period for primates, juvenile behavior in the wild is not actually well studied (Barale et al., 2015). Juvenile primates do not gain independence from their mothers until they are behaviorally and physically capable of dealing with the demands of their environment (Nowell & Fletcher, 2007). Apes in particular have long dependency periods, with offspring not weaned for up to seven years (Van Noordwijk & Van Schaik, 2005). For western lowland gorillas, complete weaning age appears to be influenced by food availability with the development of feeding skills an important part of immature primate development (Nowell & Fletcher, 2007).

How individuals will allocate their time is dependent upon the environment inhabited, social structure and food availability (Dunbar et al., 2009) and individuals must learn how to balance their time appropriately as spending more time in certain activities will be done at the expense of other activities (Hamel & Côté, 2008). In addition, time allocation can vary not only based on age and body size but also depending on sex and rank (Canteloup et al., 2019; Kurihara, 2016). Juveniles also differ from adults in terms of their behaviors; for example, juveniles are often found to rest less (Agetsuma, 2001; Prates & Bicca-Marques, 2008), play more (Barale et al., 2015; Liao et al., 2018; Masi et al., 2009; Shanee & Shanee, 2011) and spend more time feeding than adult males (Masi et al., 2009; Rothman et al., 2008).

Thus, the juvenile phase is crucial for individuals to not only learn later-life social rules, but also about the food available to them and where to find it. In many warm-blooded mammals there is a close relationship between body size and dietary choice with smaller animals often feeding on smaller, high-quality foods while larger animals will feed on greater volumes of lower-quality food (Agetsuma, 2001). For example, in Japanese macaques adult males spend more time consuming lower-quality, abundant foods compared to immatures who targeted less abundant but higher quality food sources (Agetsuma, 2001). In addition, the skill level of an individual will also influence the types of foods eaten; food sources such as fruits often require particular skills to maximize efficiency during consumption and with minimal waste (Nowell & Fletcher, 2008).

While sex differences in diet and time budgets are well studied in adults (e.g. Almeling et al., 2017; Isbell & Young, 1993; Liu et al., 2016; Rose, 1994; Shanee & Shanee, 2011; Sueur et al., 2011), work remains to be done on how age and sex influence juvenile behavior and the age at which adult-like differences become apparent. If immature individuals are using the period prior to maturity to develop the skills for adulthood, it would be expected that sex differences would be apparent during this time (Barale et al., 2015) as males and females may require different skills for success. Indeed, Black-and-gold howler monkeys (*Alouatta caraya*) spent varying amounts of time in different behaviors with the influence of sex being age dependent (Pavé et al., 2016). For species living in female-bonded groups, such as chacma baboons (*Papio ursinus*) female-female relationships are a central part of the group structure, influencing the health and survival of individuals and their offspring (Silk et al., 2003, 2006). Therefore, juvenile females may start to invest in these relationships at a young age, while this may be of less importance for males, who often disperse from their natal group (Alberts & Altmann, 1995). For species where both sexes migrate from their natal group, such as western lowland gorillas (Forcina et al., 2019), the differences in activity budget as a result of reproductive strategies may be less obvious.

Western lowland gorillas inhabit the lowland forests of Angola, the Central African Republic, Gabon, Equatorial Guinea, Cameroon and Republic of Congo (Maisels et al., 2018), and are the largest extant primate (Masi et al., 2009). Listed as Critically Endangered by the IUCN Red List (Maisels et al., 2018), habitat destruction, hunting and disease pose continued risks to western lowland gorillas (Caillaud et al., 2006; Fünfstück & Vigilant, 2015; Genton et al., 2012), making it ever more important to improve our understandings of their behavior and diet. Western lowland gorillas live in groups ranging from one mated pair to 22 individuals, typically with one silverback male, adult females (sex ratio estimated to be 0-9 females per adult male), subadults, juveniles and infants (Magliocca et al., 1999). Both male and female offspring disperse once maturity is reached (Forcina et al., 2019), with secondary transfers observed in females (Stokes et al., 2003). Immature individuals are a key part of the social unit and appear to be central to facilitating social interactions during intergroup encounters (Forcina et al., 2019).

Wild western lowland gorillas have become more central to research in recent years but much remains to be learnt about this species in their natural environments. Difficulties habituating groups, given the dense environments inhabited and shy nature of these gorillas make behavior research challenging (Hagemann et al., 2018; Shutt et al., 2014). Few studies have been conducted on wild habituated groups through full-day observations (e.g. Masi et al., 2009) while the majority of research has occurred at ‘bais’, open swampy areas within the forest where gorilla groups come to forage (e.g. Breuer et al., 2007, 2009; Klailova et al., 2010; Nowell & Fletcher, 2007, 2008; Parnell, 2002; Remis, 1997; Stokes, 2004). These have provided helpful insights into western lowland gorilla behavior, diet and sociality, however some estimates suggest that some groups can spend as little 1% of their time at the bais (Hagemann et al., 2018) meaning many important interactions and behaviors are missed.

Gorillas are sexually dimorphic, with adult males being around 75kg heavier than females (Leigh & Shea, 1995). As this dimorphism develops, the way in which individuals meet their energetic requirements may vary and influence the types of foods eaten, although adult male mountain gorillas have been found to spend the same amount of time feeding as females and juveniles (Rothman et al., 2008). However, to meet their higher energy demands, adult males eat faster to consume larger quantities of food (Rothman et al., 2008). Whether or not the same is true for western lowland gorillas is currently unknown. Masi and colleagues (2009) found that the subadult male spent more time feeding than juveniles while the adult male spent less time feeding than both adult females and immature individuals. Immature individuals were also found to spend more time in social or other activities than adult females, however it remains unknown at what age these differences become apparent and when sex begins to influence behavioral differences.

Here we study the behavior and food-part choices of a reintroduced group of immature western lowland gorillas. Using reintroduced gorillas provides a rare opportunity to observe immature western lowland gorillas in their natural habitat. In addition, our study will also contribute to a better understanding of the behavioral and dietary diversity shown by reintroduced individuals. We predict that (1) (i) as gorillas age, they will allocate their time differently due to physiological and behavioral differences with less time spent playing and more time feeding to compensate for an increasing body size; and (ii) sex differences in behavior will become apparent in older individuals as a result of differences in characteristics and skills required for adulthood with (older) males spending more time playing to build up social dominance skills and more time eating to compensate for larger body sizes as they develop. Furthermore, we predict that (2) (i) age groups will differ in the plant parts eaten as differences in body size, strength and skills will influence the ability of individuals to access different food types with younger individuals consuming less roots but more of accessible food items such as flower and leaves; and (ii) there will be no difference in the plant parts consumed between males and females as reproductive demands will not yet influence nutritional requirements. Finally, we predict that (3) (i) behavioral diversity will vary dependent on age classes, with younger individuals showing more diverse behavior ranges as they learn about their environments; (ii) while no variation in behavioral diversity will be found between the sexes as reproductive costs will not yet influence behavior; and (4) (i) dietary diversity will be influenced by age classes and younger individuals will have more diverse diets as they are less constrained by the volume of food required to meet daily needs and as they learn about appropriate foods to consume, while (ii) no sex differences will be apparent in dietary diversity as reproductive nutritional demands will not yet influence diet. Behavioral diversity has not previously been investigated for this species, but has been shown to be an important variable in enhancing conservation policies (Brakes et al., 2019) and as a potential positive indicator of welfare (Miller et al., 2020). Here we provide a first baseline for behavioral diversity in immature western lowland gorillas, which may assist future reintroductions of young gorillas by ensuring individuals are showing age appropriate behaviors.

## Methods

### Study population

We collected on western lowland gorilla (*Gorilla gorilla gorilla*) orphans who underwent a rehabilitation program and were released in the Republic of Congo (hereafter Congo) in June, 2001. Research was conducted in the Réserve Naturelle des Gorilles de Lesio-Louna (-3°28’S, 15°48’E). Elevation ranges from 300m – 750m in the reserve (King, 2011). The site is characterized by *Hyparrhenia* and *Loudetia* grasslands with swampy gallery forests along the watercourses (King et al., 2003). Average annual rainfall is estimated to be 1660mm with the main rainy seasons from October - November and March - April, the dry season from late May – September and a drier period in January and February (King, 2011).

The study group consisted of nine individuals, eight of whom were rescued from the illegal pet trade in Congo and one individual who was born to parents undergoing the rehabilitation process, but was subsequently rejected by his mother and hand-raised (King et al., 2012). The group was composed of four males and five females, ranging from 3.5 – 7.75 (mean ± SD: 5.5 ± 1.48) years old (Table 1) at the beginning of the observation period. The age of the eight rescued individuals were estimates based on their physical size, teeth and any pre-arrival information (King et al., 2012). From June 2001, the group were nutritionally independent and lived in complete freedom in the reserve whilst being monitored almost daily by local staff (King et al., 2003).

**Table 1:**
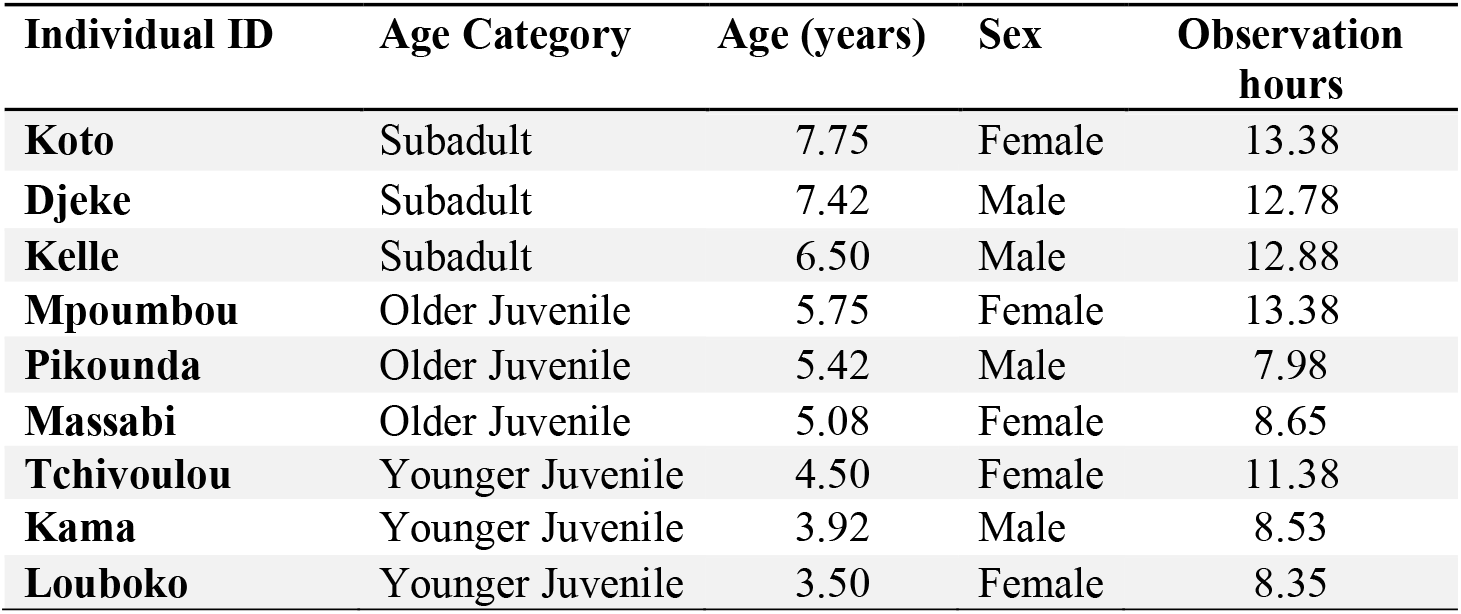
Composition of the study group and observation hours for each western lowland gorilla observed in Réserve Naturelle des Gorilles de Lesio-Louna, Republic of Congo, from September to December 2002.

### Data collection

We collected data over 25 days from the 8^th^ of September to 6^th^ of December 2002, with a mean observation time per gorilla of 10.91 hours (SD = 2.51, range = 7.89 – 13.83). All individuals in the study group were observed in September and October, while in November only six of the individuals (Djeke, Kelle, Koto, Massabi, Mpoumbou and Tchivoulou) were observed and only three individuals (Djeke, Kama, Kelle) were observed in December. We divided each day into four time periods (08:30 – 10:59; 11:00 – 12:59; 13:00 – 15:29; 15:30 – 18:00). We collected data on one or two individuals per day, with observations conducted in up to three time periods a day. The sample individuals were chosen randomly prior to finding the group each day.

We recorded data through focal animal sampling (Altmann, 1974) with instantaneous data collected at 1-minute intervals over a 50-minute period. At each minute interval, the focal animal’s activity and if feeding, the food type (if known) and food part were recorded. Continuous data were collected on target behaviors throughout the 50-minutes. Target behaviors were those which occurred less frequently or as part of a behavior sequence and allowed rarer or shorter length behaviors to be recorded (King et al., 2003). A total of 69 different behaviors were recorded; full details and descriptions of the behaviors can be found in Supplementary Material 2. The use of two separate recording methods allowed for the collection of data on the wide range of behaviors displayed by gorillas (King et al., 2003). We conducted an inter-observer reliability study in August 2002, prior to data collection and generally high rates of consistency were found (see King et al., 2003).

### Ethical Statement

Field data used in this research were collected in 2002 under the authorization of the Ministère de l’Economie Forestière of the Republic of Congo, and the Aspinall Foundation, U.K. This research adhered to the International Primatological Society Code of Best Practice Guidelines for Field Primatology. The research presented in this paper was authorized by the Aspinall Foundation, U.K.

#### Data availability

Data used in this analysis are available upon request.

### Statistical analysis

Following the age groups used in other studies of western lowland gorillas (e.g. Magliocca et al., 1999; Parnell, 2002), we divided individuals into three age categories: subadult (aged 6-8 years); older juvenile (aged 5-6 years) and younger juvenile (aged 3-5 years). Western lowland gorillas under the age of four years are often considered infants as they are still dependent on their mothers (see Breuer et al., 2009 for summary), however as our individuals were orphans and all weaned, we felt even the youngest would best fit into the juvenile age category.

#### Activity budget

To estimate the activity budget of the gorillas, we categorized the instantaneous behaviors into seven categories: eat, locomotion, rest, self-play, social play, social interaction and other (details of behaviors in Table 2). Other was categorized as behavior that did not fit into the previous categories. Due to the age range of the study group, we expected play to be an important part of their activity budget; social play and self-play were therefore not combined as previous studies have found differences in the type of play among young gorillas, with rates of this being influenced by age and sex (Maestripieri & Ross, 2004). We calculated the activity budget by combining the total counts of each behavior and dividing this by the total number of occurrences of all behaviors to give the percentage for each category.

**Table 2:**
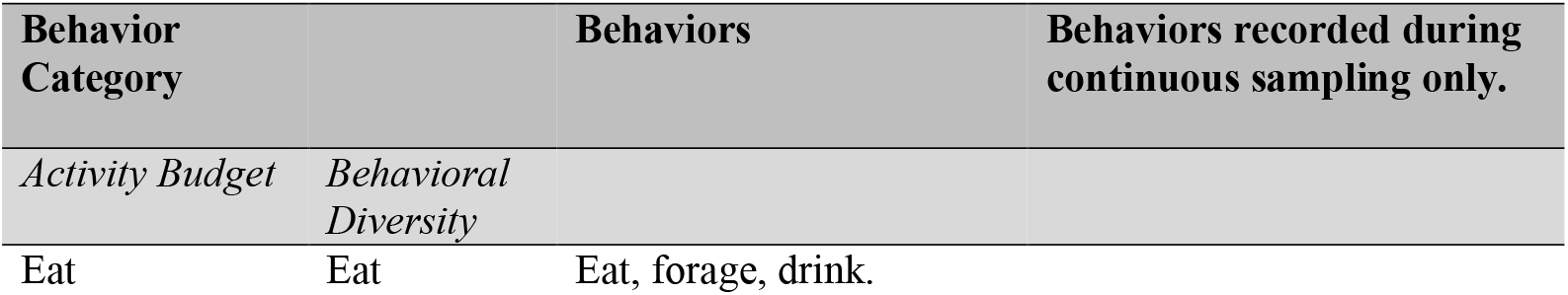
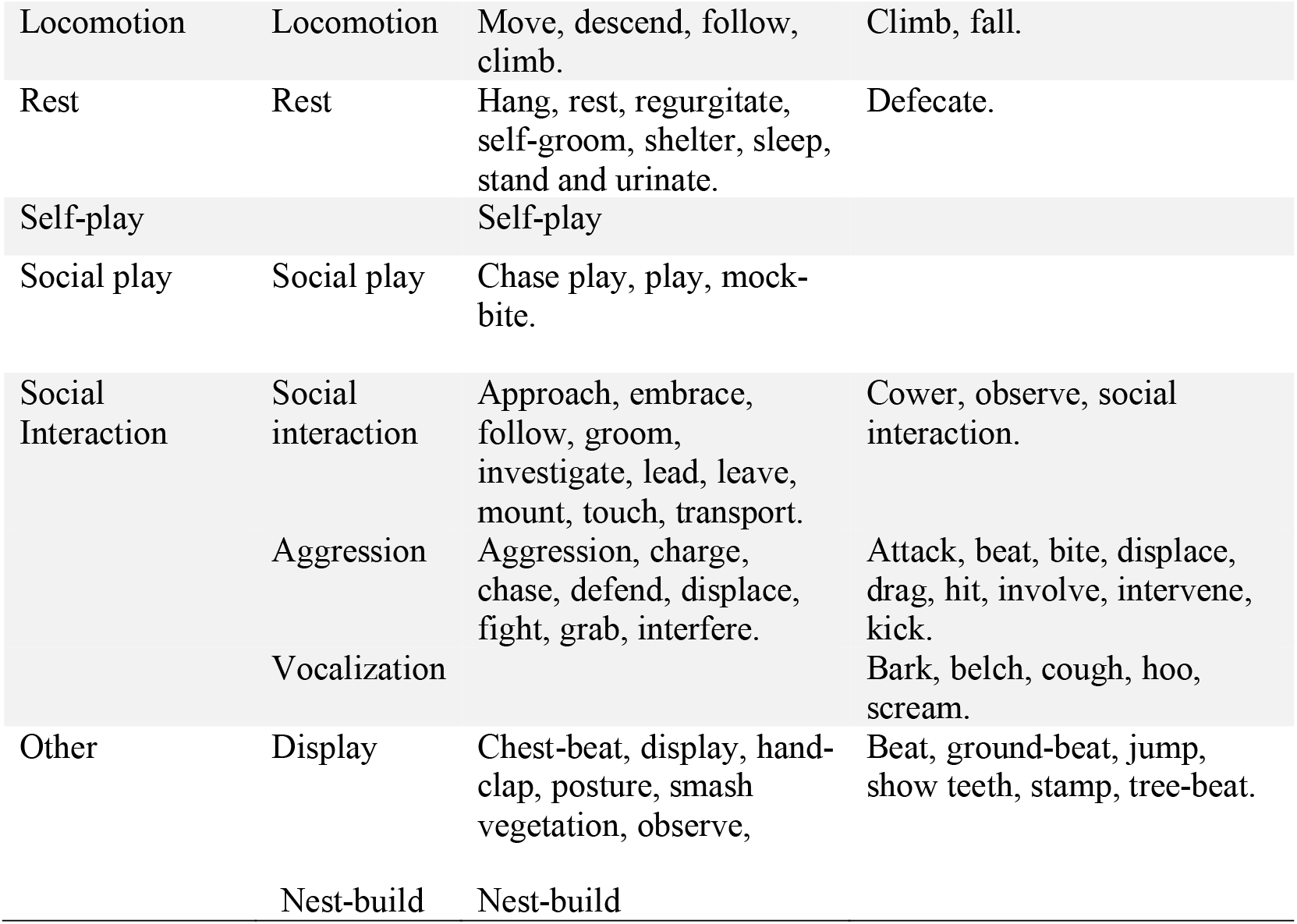
Details of western lowland gorilla behaviors used in analysis from a study population living in Réserve Naturelle des Gorilles de Lesio-Louna, Republic of Congo, observed from September to December 2002. The full ethogram can be found in Supplementary Material 2.

To analyze if there were any differences in behavior dependent on the age or sex of the study group, we ran seven Generalized Linear Mixed Methods Models (GLMMs) using a Poisson error structure. For each model, we fitted the counts of occurrence of each behavior category per time period as the response variable and fitted sex and age category as fixed effects. We included two random effects in these models: individual ID to prevent pseudoreplication and time period (four levels) to account for the differences in observation time between individuals, fitting each model with an offset of the total amount of observation of each individual in the different time periods (as data were instantaneous the total observation time was the counts of occurrence of all behaviors). The GLMM for social interaction produced skewed results; to rectify this, we transformed the dependent data and used a linear mixed model with a Gaussian error structure. We ran each model twice, changing the reference category to allow comparisons to be made between all three age categories.

#### Dietary composition

We calculated food part preference similarly to activity budget, dividing the total counts of each food part by the total count of all food parts and multiplying by 100, providing a percentage for each food part. To examine how age and sex influenced dietary composition, we divided food parts into five categories: flowers/leaves, fruits, roots, stem and other (feces, insects, other plant parts, milk). We chose food part over plant species as the records on species were incomplete due to uncertainties in the plant species during data collection. We ran five GLMMS (one per food part category), and used the total number of each food part consumed per month per individual as the response variable in the model. We again fitted each model with an offset to account for the difference in observation hours between individuals, using the count of occurrences of all food parts per month per individual. We fitted age category and sex as fixed effect and used a Poisson error structure. To prevent pseudoreplication and to account for seasonality, we fitted individual ID and month as the random effects. Month was chosen in these models rather than time period as some seasonal variation in food availability could be expected. As our study took place over a relatively short period, we did not expect there to be large variation but it was important to control for this aspect of variation. As with our behavior models, we ran each model twice, changing the reference category to allow comparisons between all age categories.

#### Behavioral and dietary diversity

To examine differences in behavioral and dietary diversity among the study group, we calculated Shannon’s Diversity Indices (SDI). Using the following equation:

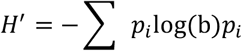

where p_i_ is the proportional abundance of a behavior/food part and b is the base of the logarithm. A larger H’ value indicates greater diversity and the value will depend partly on the number of behaviors in the sample and partly on the distribution of those behaviors (Henzi et al., 1997; Shepherdson et al., 1993). The SDI is one of the most common measures for looking at behavioral diversity (Miller et al., 2020). The higher the diversity score, the more diversity is present (Nsekanabo et al., 2022). If the SDI equals 0, there is no diversity, for example, the study population eating only one food type.

We used narrower behavioral categories than in the above analysis to allow for more precision in the diversity scores and differentiation between behaviors (Table 2) and food part choice (flower, leaves, fruit, feces, insect, root, stem, other [other plant part, milk]). To calculate behavioral diversity, we combined the total number of occurrences of each behavior from both instantaneous and continuous observations and these were used to calculate the SDI for each individual. To calculate dietary diversity, the counts of food part recorded during each focal observation were combined for each individual and the SDI calculated for each focal observation. These SDI’s were then combined for each individual and the mean SDI calculated. We did not calculate the mean SDI for behavioral diversity as the use of continuous and instantaneous data meant that this was not possible. We reasoned it was better to include the continuous data as it allowed for the inclusion of rarer or shorter behaviors. We used a Mann-Whitney test to test for differences between behavioral diversity scores and sex and Kruskal-Wallis tests to test for differences in diversity scores between age categories.

#### Statistical Tests

All statistical tests were conducted in R version 4.0.2 (R Core Team, 2020). We ran our statistical models presented above using the lme4 (Bates et al., 2015) and lmerTest (Kunzetsova et al., 2017) packages and we computed the SDI using the vegan (v. 2.5-7, Oksansen et al., 2020) package. We checked model fits using QQ plots and histograms and all models were checked for overdispersion using the DHARMa package (v. 0.4.1, Hartig, 2021). When models were singular, we checked the robustness of the results using the blme package (Chung et al., 2013). P-values were considered significant when ≤0.05. The conditional R^2^ Trigamma (r.squaredGLMM function) from the package MuMIn (v 1.43.17, Barton, 2020) was calculated for each behavior to assess the model fit and allow comparisons across studies (Harrison et al., 2018; Nakagawa & Schielzeth, 2013).

## Results

### Activity budget

The gorillas spent most of their time eating (37.5%, n=2211/5894), followed by resting (31.85%, n=1877/5894), locomotion (12.06%, n=711/5894), social play (9.76%, n=575/5894), self-play (3.61%, n=213/5894) and other (3.04%, n=179/5894), with the least amount of time spent in social interactions (2.17%, n=128/5894). Out of the seven behavior categories, we found age class differences in three of them: younger individuals spent significantly more time in locomotion than subadults (*β* ± s.e. = 0.326 ± 0.160, x^2^ = 2.037, p = 0.0416) and significantly more time in self-play compared with older juveniles (*β* ± s.e. = 1.225 ± 0.421, x^2^ = 2.909, p = 0.00363) (Figure 1). Younger juveniles also spent significantly less time resting compared with both older juveniles (*β* ± s.e. = -0.257 ± 0.113, x^2^ = -2.268, p= -0.0233) and subadults (*β* ± s.e. = -0.399 ± 0.115, x^2^ = -3.473, p = 0.000515) (Figure 1). There were no statistically significant differences between age categories in the time spent eating, social playing, in social interactions or other behavior (full results available in Supplementary Material 1; Table 3 & 4). No sex differences were found in immature gorillas activity budgets (Figure 2) (Supplementary Material 1; Table 3).

**Figure 1.**
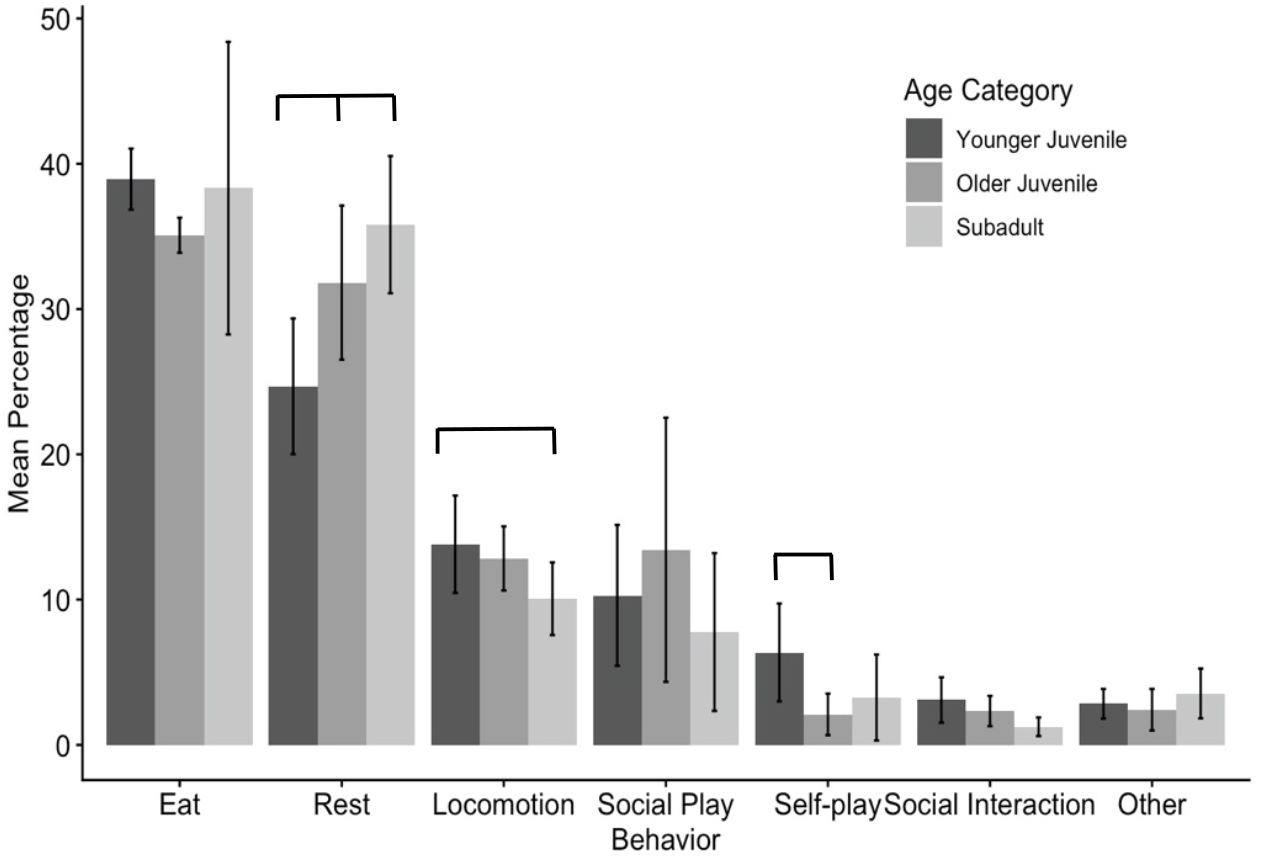
Percentage of time spent in each behavior by age group. Error bars represent standard deviation (±) from the mean. Results found to be significant in GLMMs are indicated with grey lines (younger juvenile rest when compared with older juvenile and subadults, younger juvenile locomotion when compared with subadults and self-play in younger juveniles when compared with older juveniles).

**Figure 2.**
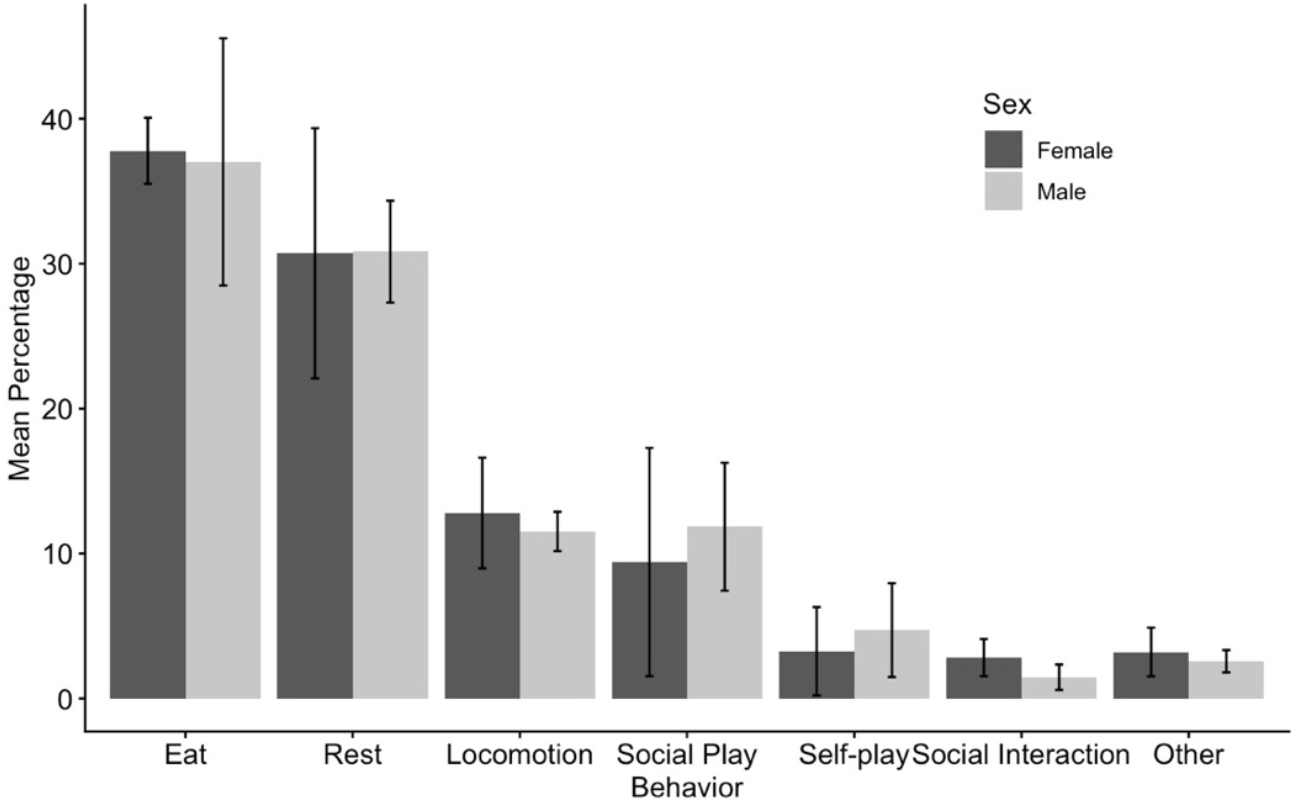
Mean percentage of time spent in each behavior by sex for a group of western lowland gorilla observed in Réserve Naturelle des Gorilles de Lesio-Louna, Republic of Congo, from September to December 2002. Error bars represent standard deviation (±) from the mean.

### Dietary composition

The gorillas fed mostly on stems (26.62%, n=541/2032), flower/leaves (24.90%, n=506/2032) and fruit (24.31%, n=494/2032), while roots and other food parts were consumed least (5.95% (n=121/2032) and 18.21% (n=370/2032) respectively). Subadults ate significantly more stems than both older and younger juveniles (*β* ± s.e. = -0.854 ± 0.381, x^2^=-2.242, p = 0.0249) (Figure 3). Males ate significantly more roots than females (*β* ± s.e. = -0.525 ± 0.235, x^2^=-2.234, p = 0.0244) (Figure 4). No other significant effect of age or sex was found on food part consumption (full results available in Supplementary Material 1; Table 5).

**Figure 3.**
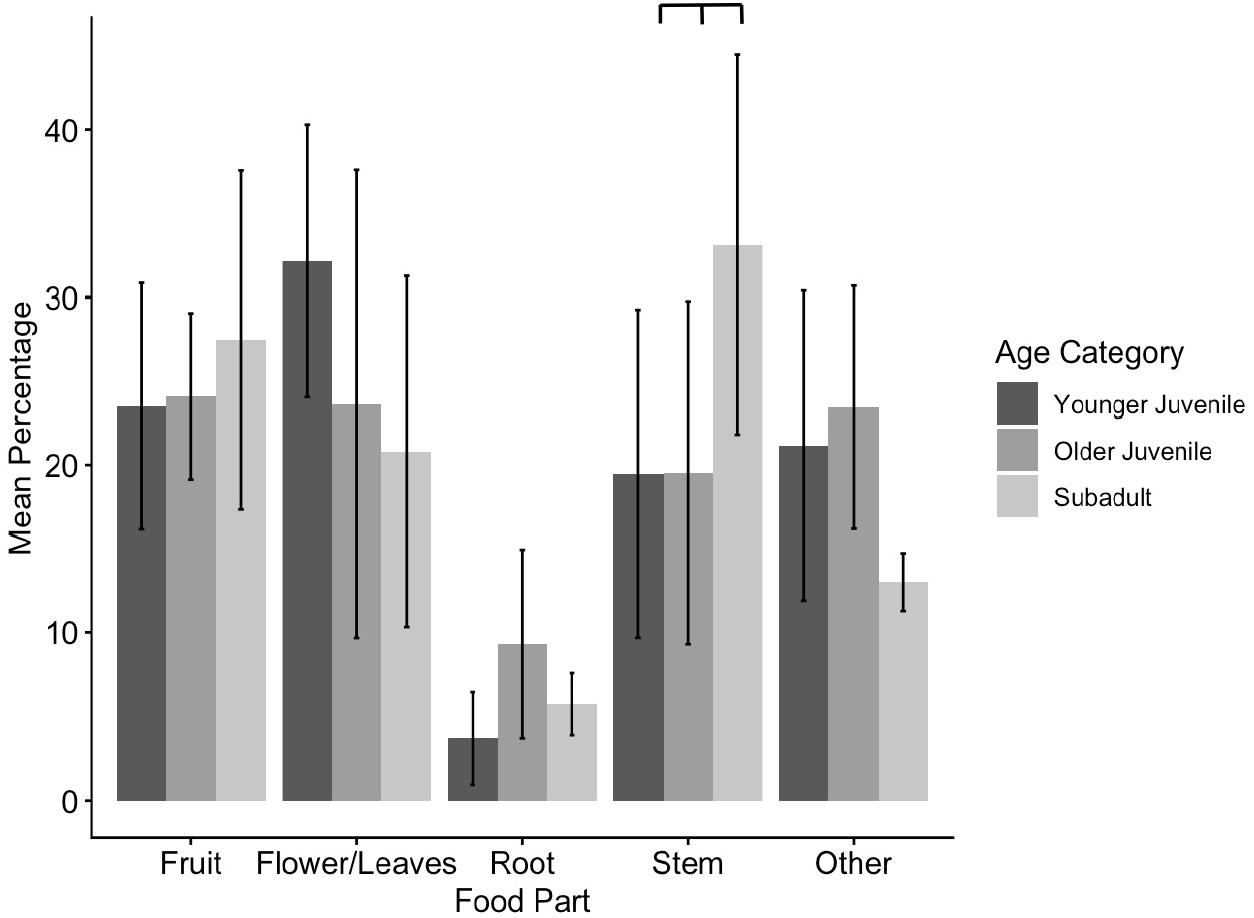
Mean percentage of diet composition by age group of western lowland gorillas observed in Réserve Naturelle des Gorilles de Lesio-Louna, Republic of Congo, from September to December 2002. Error bars represent standard deviation (±) from the mean. Results found to be significant in GLMMs are indicated with grey lines.

**Figure 4.**
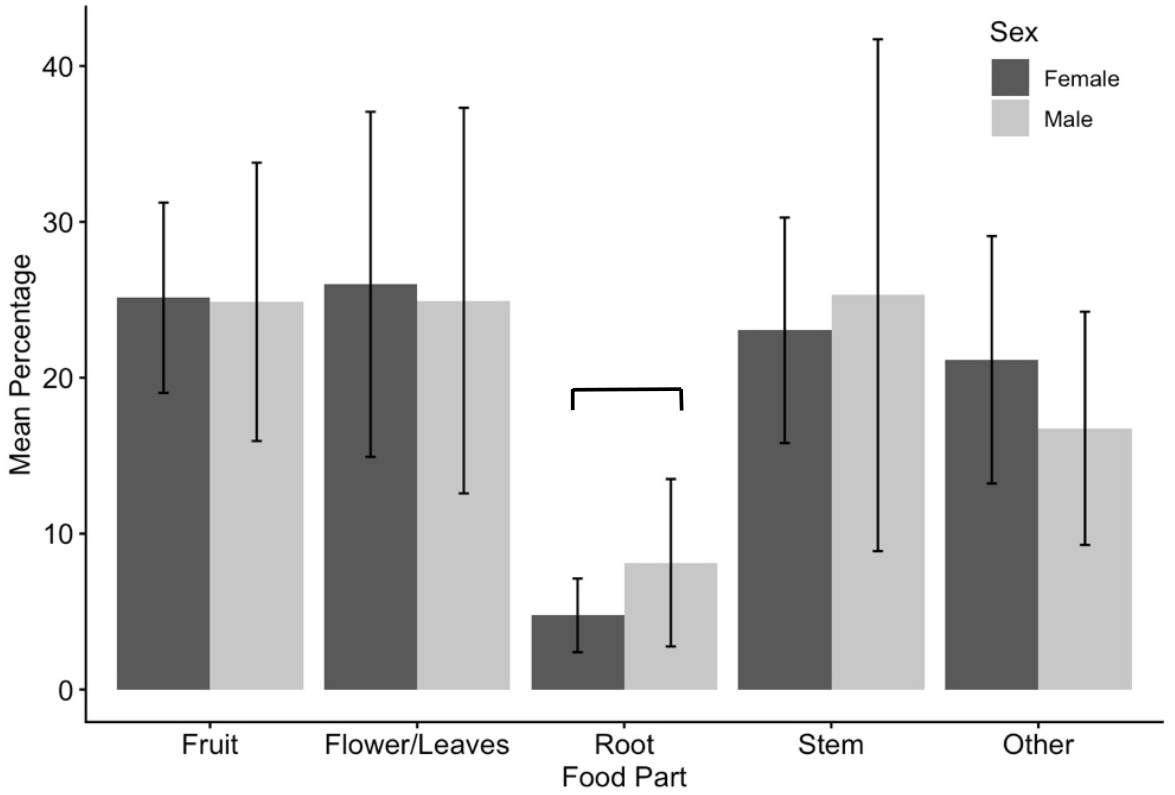
Mean percentage of diet composition by sex of a group of western lowland gorilla observed in Réserve Naturelle des Gorilles de Lesio-Louna, Republic of Congo, from September to December 2002. Error bars represent standard deviation (±) from the mean. Results found to be significant in GLMMs are indicated with grey lines.

### Behavioral and dietary diversity

Individual behavioral diversity values ranged from 1.640 – 2.008 (mean ± SD = 1.844 ± 0.120, n=9). Neither sex nor age affected individual behavioral diversity significantly (sex: W = 5, p-value = 0.286; age: chi-squared = 0.356, df= 2, p=value = 0.837, n = 9). Dietary diversity ranged from 0.458 – 0.786 (mean ± SD = 0.664 ± 0.109, n = 9). No age or sex differences were found (sex: W = 9, p = 0.905, age: chi-squared = 3.822, df = 2, p-value = 0.148, n = 9).

## Discussion

As western lowland gorilla populations decrease, it is ever more important to understand their behavioral ecology to aid their protection and conservation. While we found some age-class differences in immature gorilla behavior and food choices, there were no differences between the sexes in any of our variables, suggesting that the sexes do not begin to diverge during the juvenile stage. The results provide insights into immature gorilla behavior and can help to form a basis for future studies; especially the data on behavioral diversity are valuable for future reintroductions. Reintroduction biology is still a relatively new and developing area (Seddon et al., 2007) and therefore providing some baseline data for species will be highly beneficial.

### Activity Budget

Individuals have a finite amount of time to dedicate to their daily activities (Dunbar et al., 2009) and more time spent in one activity will result in less time for other behaviors. Different developmental stages and energetic requirements dictate how individuals allocate their time dependent on their age and sex. The activity budgets for each behavior in our study were within similar ranges to those observed in a study by Le Flohic and colleagues (2015), who similarly studied a group of rehabilitated immature western lowland gorillas in their natural habitat. However, Masi and colleagues (2009) reported significantly higher feeding (61.1% - 76.1% dependent on season) and lower resting times (21%). The key difference between the study groups was the artificial composition of our study group and that of Le Flohic and colleagues (2015), both composed of immature orphaned individuals, while the group Masi et al. (2009) studied was naturally composed with adults, a subadult, juveniles and infants. The lack of adults may have been a key factor in influencing the differences in activity budget between study populations (Le Flohic et al., 2015), as could the lack of mothers to learn from (Megna, 2002). Human influences on the group during the lengthy pre-release rehabilitation phase may also have been a factor.

Although it is well known that group size affects activity patterns (Lehmann et al., 2008), group sizes were comparable across these three studies, ranging from 5-13 individuals with our study group in the middle. Interestingly, the two reintroduced groups of immature individuals showed very similar activity patterns. Another factor affecting activity budgets is habitat ecology (Canteloup et al., 2019; Dunbar et al., 2009; Hanya, 2004). Both our study and Le Flohic and colleagues (2015) occurred in the Batéké Plateaux (in Congo and Gabon respectively), an area characterized by grasslands and swampy gallery forests, while Masi and colleagues carried out research in the Central African Republic, at Bai Hokou; an area of continuous forest. Rainfall is also greater in the Batéké Plateaux (1660mm average in the Lesio-Louna Reserve; King, 2011) compared with 1365mm average in Bao Hakou (Remis et al., 2001). These variables might further explain differences in feeding and resting times found between groups. The greater proportion of time spent resting in our study suggests that individuals were able to meet their energetic requirements more quickly, which might be due to a lack of food competition. Alternatively, it could be that our individuals required more time for digestion due to lower quality food being consumed.

We found that younger juveniles spent more time in self-play and resting compared with older juveniles. Play behavior has previously been shown to decrease when individuals get older (Shimada & Sueur, 2018) and as play behavior usually only occurs when other needs are met (Boissy et al., 2007), the decreased time spent resting and increased time spent in self-play suggests that younger juveniles require less time to rest and are able to meet their daily needs more quickly, freeing time for other activities, such as self-play. Higher levels of self-play among younger juveniles may reflect the need for skill development and learning of non-social skills (Montgomery, 2014). Our results are also in line with those of a study of captive western lowland gorillas, where self-play decreased from three years onwards (Maestripieri & Ross, 2004). The difference in time spent resting may be a result of body size, with heavier individuals spending more time resting compared with lighter individuals (Agetsuma, 2001; Masi et al., 2009).

Interestingly, our prediction that there would be sex differences in activity budgets, especially in older individuals, was not supported. This suggests that sex differences in the activity budget do not become apparent until later in life. Studies of mountain gorillas found that males and females do not significantly differ in body measurement categories until ages 12-18 years old with initial size difference not emerging in body size until 8.5-10 years old (Galbany et al., 2017). Unfortunately, our sample size was too small to assess if sex differences in behavior were starting to emerge in older juveniles only. For western lowland gorillas sex differences in activity budget may become apparent even later in life, as compared to mountain gorillas they are weaned later (Breuer et al., 2009).

### Dietary composition

The diet of western lowland gorillas is highly seasonal, especially when compared with relatively stable diet of mountain gorillas (Nowell & Fletcher, 2008), therefore individuals must learn to access a variety of food sources easily to be able to adapt to the changing food availability. We found stem consumption was higher among subadults compared to younger individuals. Two reasons are suggested for this; one, more strength or skills may be required to access stems as a food source with body size and skills influencing food consumption. For younger individuals to access this resource, it may require more time or skills and preference is placed to other food items. Alternatively, due to the larger size of subadults, they may need to utilize alternative food sources to meet their daily requirements. In line with this, mountain gorilla silverbacks eat more per day than females and juveniles, but do not spend more time eating, suggesting they utilized different food sources to meet their energetic needs (Rothman et al., 2008). Young individuals must find the balance between time spent processing foods and feeding to optimize the benefits available from each food resources (Nowell & Fletcher, 2008).

It could be expected that once gorillas have reached adulthood, the significant dimorphism between adults may influence food choices as the considerably larger size of adult males may affect access to food sources. However, results from a study of western lowland gorilla diet across sites did not indicate a sex difference in diet (Doran et al., 2002). As our study group was composed of males and females, we did not expect to see an overall sex difference in food part choice. Our finding that males ate more roots than females was thus unexpected and may be a result of personal preferences, different nutritional requirements or strength levels. Individual experiences can influence foraging specialization (Durell, 2000) and as the individuals in our study group all originated from different backgrounds, this too may have had an influence on their food preferences. As the species of plants consumed were unknown, we are unable to determine whether this was roots across a number of plants or one species in particular that led to the increased consumption among males.

### Behavioral and dietary diversity and reintroduction success

Studying behavioral and dietary diversity allows the entire behavior repertoire and diet composition to be examined. Knowing behavioral diversity has important conservation implications and can be used to measure whether conservation efforts are assisting or restricting a population. Chimpanzees (*Pan troglodytes ellioti; Pan t. schweinfurthii, Pan t. troglodytes, Pan t. versus*) living in areas with high levels of human disturbance had lower behavioral diversity compared with those living in areas with lower levels of disturbance (Kühl et al., 2019). Similar results were found with black howler monkeys (*Alouatta pigra*) where biodiversity loss was accompanied by behavioral diversity loss (Negrín et al., 2016). While previous studies of reintroduced groups of western lowland gorillas have looked at survival rates, reproduction success, population viability and the initial post-release period (King et al., 2006, 2009; King et al., 2012, 2014; King & Courage, 2007; Le Flohic et al., 2015), research into the behavior of reintroduced individuals is limited. The release of our study group was part of a long-term reintroduction which was considered a success, with high survival rates and multiple successful reproductions (King et al., 2012, 2014). Seven of the nine gorillas in our study group were still alive and monitored over twenty years after their release (T. King, unpubl. data). Knowing behavioral and dietary diversity scores of this group can assist future releases by providing what can be considered a normal range of diversity for post-release success. To our knowledge, it has not yet been calculated for western lowland gorillas across the whole behavior repertoire in captivity (Nsekanabo et al., 2022 examined mobility/immobility) or in the wild and by calculating both behavioral and dietary diversity, we hope to provide a baseline level which can be utilized to assist future reintroductions and be built on in future research. Additionally, the behavioral and dietary diversity levels of a free-ranging population can also be used to assist zoo practitioners in better understanding the diversity levels that can be aimed for in captive populations.

In line with our prediction, we did not find any sex related differences in diversity indices, however, contrary to our prediction we did not find any age-related differences. The environment in which a population lives may have a stronger influence on the behavioral diversity as more complex, variable environments, both in captivity and the wild have been found to illicit higher rates of behavioral diversity (Kalan et al., 2020; Scott & LaDue, 2019) while dietary diversity may be more dependent on group variability than individual parameters especially as individual foraging strategies can also be influenced by other members of the group (Durell, 2000). The small sample size and limited age range of our study group may have meant that behavioral diversity differences are less visible than in a group with a larger sample size and wider age range. For example, the behavior of dependent infants would be expected to be different to juveniles or adults as they do not require time to search for and process food. Immature western lowland gorillas use observation and imitation to develop their foraging strategies (Byrne & Byrne, 1993) and it is thus likely that feeding strategies of group members will have influenced each other. Group-living provides opportunities for social learning (Sheppard et al., 2018) and this may be reflected in both the behavior and diet of a group.

Considering diversity provides practitioners with an alternative method for measuring welfare; individuals living in stimulating environments tend to have higher behavioral diversity scores (Hall et al., 2021). Our behavioral diversity results were within a similar range to a study on captive chimpanzees (Hall et al., 2021). However, in this study, Hall and colleagues (2021) highlight the need for standardization of behaviors for future research in order to allow comparisons.

### Conclusions

Throughout western lowland gorilla research, as with many other primate species, sample sizes remain small due to difficulties habituating groups. Where studies do have larger numbers (e.g. Magliocca et al., 1999), they are conducted at bais where groups can gather during feeding. These are useful insights into gorilla behavior, ecology and structure but are restricted to a specific environment. Whilst it remains difficult to increase sample size, research using small numbers of individuals provides insights into gorilla behavior and can provide foundations for future work. Our study contributes to understanding immature western lowland gorilla behavior and provides a baseline information for the normal behavior and dietary range of rehabilitated and reintroduced gorillas. As western lowland gorillas remain under threat, further reintroductions may be needed in the future.

## Data Availability Statement

The datasets generated during or analyzed during the current study are available upon reasonable request from the corresponding author.

## Author contributions

TK developed the fieldwork methodology and supervised data collection, NN, EB and SM-R conducted the fieldwork and collected data, AC formulated the data analysis methodology with guidance from JL and TK. AC analyzed the data. AC and JL wrote the manuscript with editorial input from TK.

## Acknowledgements

We thank the Ministère de l’Economie Forestière of the Republic of Congo, and The Aspinall Foundation of UK, for permission to undertake the fieldwork for this research in the Lesio-Louna Reserve. We also thank the staff of the Projet Protection des Gorilles, Republic of Congo, for their assistance during the data collection. Waterproof notebooks were generously donated by BCB International Ltd.

## Funding

No funding was received for this research.

## Conflict of interest

The authors declare that there are no conflict of interests.

## Notes

### Competing Interest Statement

The authors have declared no competing interest.

