## Supplementary Material 1 & 2 for "Effects of age and sex on diet and activities of immature reintroduced western lowland gorillas"

Table 3: Generalized Linear Mixed Model results predicting behavior dependent on sex or age category. Younger denotes younger juvenile category. Reference category: [b] older juvenile; [c] subadult.

| Behavior | Explanatory variable | Effect size | S.E. | $X^2$ | P |
|---|---|---|---|---|---|
| **Eat** | Intercept [b] | -1.010 | 0.147 | | |
| | Sex (m) | -0.030 | 0.100 | -0.296 | 0.767 |
| R2c: 0.809 | Age category (subadult) [b] | 0.082 | 0.119 | 0.688 | 0.492 |
| | Age category (younger) [b] | 0.106 | 0.116 | 0.914 | 0.361 |
| | Intercept [c] | -0.928 | 0.156 | | |
| | Age category (younger) [c] | 0.024 | 0.118 | 0.201 | 0.841 |
| **Locomotion** | Intercept [b] | -2.049 | 0.126 | | |
| | Sex (m) | 0.018 | 0.134 | 0.130 | 0.896 |
| R2c: 0.520 | Age category (subadult) [b] | 0.268 | 0.160 | -1.670 | 0.095 |
| | Age category (younger) [b] | 0.058 | 0.153 | 0.382 | 0.703 |
| | Intercept [c] | -2.317 | 0.149 | | |
| | Age category (younger) [c] | 0.326 | 0.160 | 2.037 | 0.0416 |
| **Rest** | Intercept [b] | -1.158 | 0.124 | | |
| | Sex (m) | -0.080 | 0.096 | -0.826 | 0.4086 |
| R2c: 0.829 | Age category (subadult) [b] | 0.142 | 0.112 | 1.263 | 0.2066 |
| | Age category (younger) [b] | -0.257 | 0.113 | -2.268 | -0.0233 |
| | Intercept [c] | -1.016 | 0.133 | | |
| | Age category (younger) [c] | -0.399 | 0.115 | -3.473 | 0.000515 |
| **Social Play** | Intercept [b] | -2.401 | 0.336 | | |
| | Sex (m) | 0.548 | 0.345 | 1.590 | 0.1119 |
| R2c: 0.895 | Age category (subadult) [b] | -0.703 | 0.415 | -1.695 | 0.0902 |
| | Age category (younger) [b] | -0.157 | 0.397 | -0.396 | 0.6918 |
| | Intercept [c] | -3.104 | 0.395 | | |
| | Age category (younger) [c] | 0.545 | 0.415 | 1.313 | 0.1891 |
| **Self-play** | Intercept [b] | -4.179 | 0.361 | | |
| | Sex (m) | 0.448 | 0.357 | 1.255 | 0.20945 |
| R2c: 0.844 | Age category (subadult) [b] | 0.674 | 0.445 | 1.515 | 0.12967 |
| | Age category (younger) [b] | 1.225 | 0.421 | 2.909 | 0.00362 |
| | Intercept [c] | -3.505 | 0.398 | | |
| | Age category (younger) [c] | 0.551 | 0.424 | 1.298 | 0.194 |
| **Other** | Intercept [b] | -3.696 | 0.278 | | |
| | Sex (m) | -0.287 | 0.264 | -1.089 | 0.276 |
| R2c: 0.525 | Age category (subadult) [b] | 0.455 | 0.302 | 1.507 | 0.132 |
| | Age category (younger) [b] | 0.230 | 0.309 | 0.745 | 0.456 |
| | Intercept [c] | -3.241 | 0.291 | | |
| | Age category (younger) [c] | -0.225 | 0.305 | -0.737 | 0.461 |

Table 4: Linear mixed model predicting social interaction. Younger denotes younger juvenile category. Reference category: [b] older juvenile; [c] subadult.

| Behavior | Explanatory variable | Effect size | S.E. | T | P |
|---|---|---|---|---|---|
| **Social interaction** | Intercept [b] | -3.979 | 0.450 | | |
| | Sex (m) | -0.217 | 0.505 | -0.430 | 0.688 |
| R2c: 0.534 | Age category (subadult) [b] | -0.209 | 0.612 | -0.341 | 0.748 |
| | Age category (younger) [b] | 0.362 | 0.583 | 0.620 | 0.566 |
| | Intercept [c] | -4.188 | 0.527 | | |
| | Age category (younger) [c] | 0.570 | 0.599 | 0.952 | 0.392 |

Table 5: Generalized linear mixed models predicting food part consumption. Younger denotes younger juvenile category. Reference category: [b] older juvenile; [c] subadult.

| Food part | Explanatory variable | Effect size | S.E. | $X^2$ | P |
|---|---|---|---|---|---|
| **Flower/leaves** | Intercept [b] | -2.703 | 0.257 | | |
| R2c: 0.783 | Age category (subadult) [b] | 0.111 | 0.342 | 0.323 | 0.746 |
| | Age category (younger) [b] | 0.507 | 0.330 | 1.535 | 0.125 |
| | Intercept [c] | -2.593 | 0.296 | | |
| | Age category (younger) [c] | -0.038 | 0.339 | 1.167 | 0.243 |
| **Fruit** | Intercept [b] | -2.224 | 0.390 | | |
| R2c: 0.917 | Age category (subadult) [b] | 0.140 | 0.320 | 0.437 | 0.662 |
| | Age category (younger) [b] | -0.038 | 0.314 | -0.122 | 0.903 |
| | Intercept [c] | -2.085 | 0.418 | | |
| | Age category (younger) [c] | -0.208 | 0.275 | -0.756 | 0.450 |
| **Root** | Intercept [b] | -3.719 | 0.262 | | |
| | Sex (m) | 0.525 | 0.235 | 2.234 | 0.0255 |
| R2c: 0.561 | Age category (subadult) [b] | -0.471 | 0.259 | -1.819 | 0.0689 |
| | Age category (younger) [b] | -0.579 | 0.300 | -1.927 | -0.0540 |
| | Intercept [c] | -4.189 | 0.302 | | |
| | Age category (younger) [c] | -0.108 | 0.3194 | -0.337 | 0.736 |
| **Stem** | Intercept [b] | -2.747 | 0.331 | | |
| | Sex (m) | -0.387 | 0.323 | -1.197 | 0.232 |
| R2c: 0.897 | Age category (subadult) [b] | 0.888 | 0.384 | 2.312 | 0.0208 |
| | Age category (younger) [b] | 0.034 | 0.372 | 0.092 | 0.9271 |
| | Intercept [c] | -1.860 | 0.375 | | |
| | Age category (younger) [c] | - 0.854 | 0.381 | -2.242 | 0.0249 |
| **Other** | Intercept [b] | -2.646 | 0.290 | | |
| | Sex (m) | 0.007 | 0.324 | 0.021 | 0.983 |
| R2c: 0.879 | Age category (subadult) [b] | -0.686 | 0.393 | -1.748 | 0.081 |
| | Age category (younger) [b] | -0.259 | 0.286 | -0.671 | 0.984 |
| | Intercept [c] | -3.333 | 0.334 | | |
| | Age category (younger) [c] | 0.427 | 0.324 | 0.021 | 0.9833 |

### Supplementary Material 2

Table 6: Behavior definitions recorded during sampling.

| Instantaneous sampling | Continuous sampling | Definition |
|---|---|---|
| **Rest** | | Sit or lie with eyes open while not performing other behaviours |
| **Sleep** | | Sit or lie with eyes closed while not performing other behaviours |
| **Eat** | | Chew food item |
| **Forage** | | Find or prepare food item |
| **Regurgitate** | **Regurgitate** | Chew regurgitated food |
| **Drink** | **Drink** | Intake of fluid |
| **Defecate** | **Defecate** | Act of passing faeces |
| **Urinate** | **Urinate*** | Act of passing urine |
| **Nest-build** | | Preparation of area upon which to rest |
| **Observe** | **Observe*** | Stare at another individual, object or event |
| **Self-groom** | | Pick through own hair |
| **Self-play** | | Exhibit active behaviour alone, with no other function except self-amusement |
| **Hand-clap** | **Hand-clap** | Beating of hands together |
| | **Bark** | Short loud vocalisation resembling bark of dog, often when surprised |
| **Climb** | | Ascend vegetation |
| **Descend** | | Descend from vegetation |
| | **Fall** | Fall out of tree |
| **Approach** | **Approach*** | Move directly to within 3m of a stationary individual |
| **Leave** | **Leave*** | Move away from within 3m of a stationary individual |
| **Displace** | **Displace** | Move to within 3m of another individual who consequently moves away within 3 seconds |
| **Follow** | | Move along behind another moving individual |
| **Lead** | | Move along in front of another moving individual |
| **Move** | | Change of location unrelated to actions of another individual |
| **Groom** | | Pick through hair of another individual |
| **Touch** | | Passive contact with another individual using hand |
| **Play** | | Active contact with another individual involving contented vocalisations; occasionally may be directional |
| **Embrace** | **Embrace*** | Ventral-ventral contact interaction with another individual using both arms; may occasionally be directional |
| **Transport** | | Provide complete support to a smaller individual |
| **Investigate** | **Investigate** | Sustained contact with fingers or mouth of genital region of another individual |
| **Mount** | **Mount** | Sexual intercourse with or without penetration; may be one-directional |
| **Attack** | **Attack** | Sustained one-directional aggressive interaction with physical contact |
| **Fight** | **Fight** | Sustained two-directional aggressive interaction with physical contact; direction indicates initiator |
| **Chase** | **Chase** | Aggressive following of another moving gorilla |
| | **Bite** | Instantaneous forceful bite of another individual |
| | **Cough** | Short throaty vocalisation with lips extended forward, often used as a warning |
| | **Scream** | Long high-pitched vocalisation with wide mouth, often used during aggressive encounters |

| | | |
|---|---|---|
| | **Belch*** | Extended soft throaty vocalisation |
| **Cower** | **Cower*** | Physical exhibition of unwillingness to partake in aggressive interaction without moving away |
| **Ignore** | **Ignore*** | Lack of response to aggression or display of another individual |
| **Involve** | **Involve*** | Involvement in aggressive or non-aggressive interaction between two or more other individuals, without causing interaction to cease |
| **Intervene** | **Intervene** | Involvement in aggressive interaction between two or more other individuals, causing interaction to cease |
| **Interfere** | | Involvement in non-aggressive interaction between two or more other individuals, causing interaction to cease |
| | **Hoo** | Vocalisation used as part of display sequence |
| | **Chest-beat** | Series of beats of hands on torso |
| | **Thigh-beat** | Series of beats of hands on thighs |
| | **Tree-beat** | Series of beats of hands on tree |
| | **Beat** | Series of beats of hands on an object |
| | **Ground-beat** | Series of beats of hands on ground |
| | **Stamp** | Beat of feet on ground, one foot at a time |
| | **Jump** | Simultaneous 2-footed beat on ground |
| | **Smash vegetation** | Visual display through destruction of vegetation |
| | **Kick** | Instantaneous forceful contact of another individual with a foot |
| | **Hit** | Instantaneous forceful contact of another individual with a hand |
| | **Grab** | Forceful contact with another individual involving sustained use of hand |
| | **Drag** | Movement of another individual by sustained forceful contact with a hand |
| | **Mock-bite** | Non-forceful bite of another individual |
| | **Charge** | Rapid movement over a short distance; this may or may not be directional |
| | **Flee** | Run away from another individual during an aggressive encounter |
| | **Posture** | Stationary visual display with all 4 legs straight and lips blown out |
| **Aggression** | | Aggressive behaviour without physical contact |
| **Defend** | | Aggressive behaviour without contact as response to aggression from another individual |
| **Display** | | Physical exhibition of strength, presence or mood |
| **Hang** | | Hang from an object |
| **Stand** | | Stand on hind legs |
| **Chase-play** | | Chase another individual in a playful manner. Not aggressive. |
| **Shelter** | | Seek cover from rain |

* Behavior only recorded in continuous sampling if part of a sequence of aggression, display or other selected continuous sampling behavior.
